## Supplementary material for "Orb2 enables rare-codon-enriched mRNA expression during *Drosophila* neuron differentiation": Supp figs

**SUPPLEMENTAL DATA**

**Supplemental Datasets**

**Supplemental Dataset 1. Sequences of new transgenes presented in this study.** Columns show name of transgene (from this paper or previously published[16]), coding sequence (CDS), length of CDS, descriptive factors about CDS, and number of annotated Orb2 binding sites per kilobase.

**Supplemental Dataset 2. Data for RNAi screen.** Rows labeled for targeted gene, average fluorescence intensity value, and individual fluorescence intensity values for each animal.

**Supplemental Dataset 3. RNAseq data comparing control and *orb2* RNAi WL3 brains.** Genes that were identified in the RNA sequencing experiment, along with CAI and numbers pertaining to the RNA sequencing experiment.

**Supplemental Dataset 4. Fly stocks used in this study.** Leftmost three columns lists only the stocks used during the RNAi screen. Gene, stock number (v indicating obtained from Vienna *Drosophila* Resource Center, bd indicating obtained from Bloomington *Drosophila* Stock Center) and Fbgn number listed. Other two columns lists all other stocks used in this study, with name of stock as referred to in the study and stock number (v indicating obtained from Vienna *Drosophila* Resource Center, bd indicating obtained from Bloomington *Drosophila* Stock Center) or source listed.

__Supplemental Figure Legends__

**Figure S1: Other codon-biased reporters express at low levels in neuroblast and high levels in neurons.**

**A:** GFP-common expresses highly throughout the brain. Scale bar is 20 µm. **B-D'**: GFP-rare$^b$, mchGFP100DV1 and GFP54C3' express highly in neurons, not in NBs. **B.** GFP-rare$^b$ expression in one lobe of the *Drosophila* larval brain. Scale bar is 20 µm. **C.** Schematic of codon and sequence composition of mCHGFP100DV1 reporter (Allen et al., 2022). **C'**. WL3 Larval body expression, arrows indicate testes and brain. **C"**. Expression of mCHGFP100DV1 reporter in one lobe of WL3 brain. Scale bar is 20 µm. **D.** Schematic of codon and sequence composition of GFP54C3' reporter (Allen et al., 2022). **D'**. WL3 Larval body expression, arrows indicate testes and brain. **D"**. Expression of GFP54C3' reporter in one lobe of WL3 brain. Scale bar is 20 µm. **E.** Representative image of a MARCM clone in the central WL3 brain. Scale bar is 10 µm. Green indicates GFP-rare$^b$ protein, magenta indicates Elav staining, gray indicates mCD8 RFP highlighting clone. Arrows and weak Elav staining indicate GFP-rare$^b$ negative immature neurons. Strong Elav staining indicates mature neurons in the clone. Labelled are neuroblast (NB), immature neuron (imm. Neuron) and mature neuron. **F-I.** Representative confocal images of either GAL4 driving LacZ (**F-G)** or antibodies (**H-I**) to mark cells of interest. Scale bars are 30 µm. **F.** *OK6*-GAL4 driving LacZ expression in motor neurons in the ventral nerve cord. Some are GFP-rare$^b$ negative (dashed outline), others are GFP-rare$^b$ positive (solid outline). **G**. *engrailed*-GAL4 driving LacZ expression in serotonergic neurons in the ventral nerve cord. All are GFP-rare$^b$ positive (solid outline). **H.** Wrapper antibody staining indicating midline glia in ventral nerve cord. Dashed outline indicates cell bodies without GFP-rare$^b$ protein. **I**. Repo antibody staining in ventral nerve cord area rich in astrocyte-like glia. Most are GFP-rare$^b$ negative, indicated by dashed outline. **J-K.** Quantification of Type I (J) and Type II (K) NB numbers in WL3 brains of control and *insc*-GAL4 driving UAS-RNAi against *brat, numb* and *prospero.* Each data point = 1 animal. Multiple biological replicates were performed. *=p<0.05, **=p<0.01. ****=p<0.0001 by one-way ANOVA followed by Dunnett's multiple comparisons test.

**Figure S2: Orb2 whole-animal knockout and neuron specific knockdown result in similar decreases in GFP-rare$^b$ expression. A:** Knockout of *orb2* in the entire animal leads to a more than 2-fold decrease in GFP-rare$^b$ expression. Each data point=1 animal. Multiple biological replicates performed. Unpaired t test performed, **** = p<0.0001. **B-D**: GFP-rare$^b$ protein fluorescence in WL3 brains of indicated genotypes. Scale bar = 10 µm. **E**: Quantification of GFP-rare$^b$ protein fluorescence in indicated cell types in the indicated genotypes. One-way ANOVA was performed followed by Šídák's multiple comparisons test. *: p<0.05. ****: p<0.0001. **F:** Quantification of GFP-rare$^b$ protein expression in total brain for WT (*w* RNAi), second *CG4612* (v52947) RNAi, and second *CG13928* (v51777) RNAi compared to average fluorescence intensity of WT total brains. Each data point= 1 animal, multiple biological replicates were performed. ***=p<0.005 by One-way ANOVA followed by Dunnett's multiple comparisons test**.** **G-H**: Representative confocal images of adult testes expressing GFP-common, scale bar is 50 µm. **G**: *vasa*-GAL4 expressing *w* RNAi. **H**: *vasa*-GAL4 expressing *orb2* RNAi. **I**: Quantification of reporter GFP-common expression in testes under either control (*w* RNAi) or *orb2* RNAi driven by *vasa*-GAL4. Line profiles of GFP of protein expression taken from hub to spermatid (300 µm). Welch's t test comparing linear regressions shows no difference.

**Figure S3**: **Both Orb2 binding sites and CAI are important to determine expression regulation in the *Drosophila* brain**. **A:** Graph of CAI of each codon in the sequence of the GFP-rare$^{b}$ transgene with annotated Orb2 binding sites marked in green. **B**. Graph of CAI of each codon in the sequence of the GFP-common transgene, no Orb2 binding sites to be annotated. **C-D.** Scatter plots of number of annotated Orb2 binding sites in the 3' quartile of the CDS and the 3' UTR of transcripts of interest normalized to length, plotted against CAI of each transcript. Linear regression line plotted, r and p value for shown on the graph. **C.** Transcripts that are significantly upregulated by Orb2. **D.** Transcripts that are significantly downregulated by Orb2. **E.** Selected rare-codon-enriched genes that pass screening criteria (**Fig 5E**). Fold change values displayed in green, binding sites displayed in blue.

### Figure S1: Other codon-biased reporters express at low levels in neuroblast and high levels in neurons.

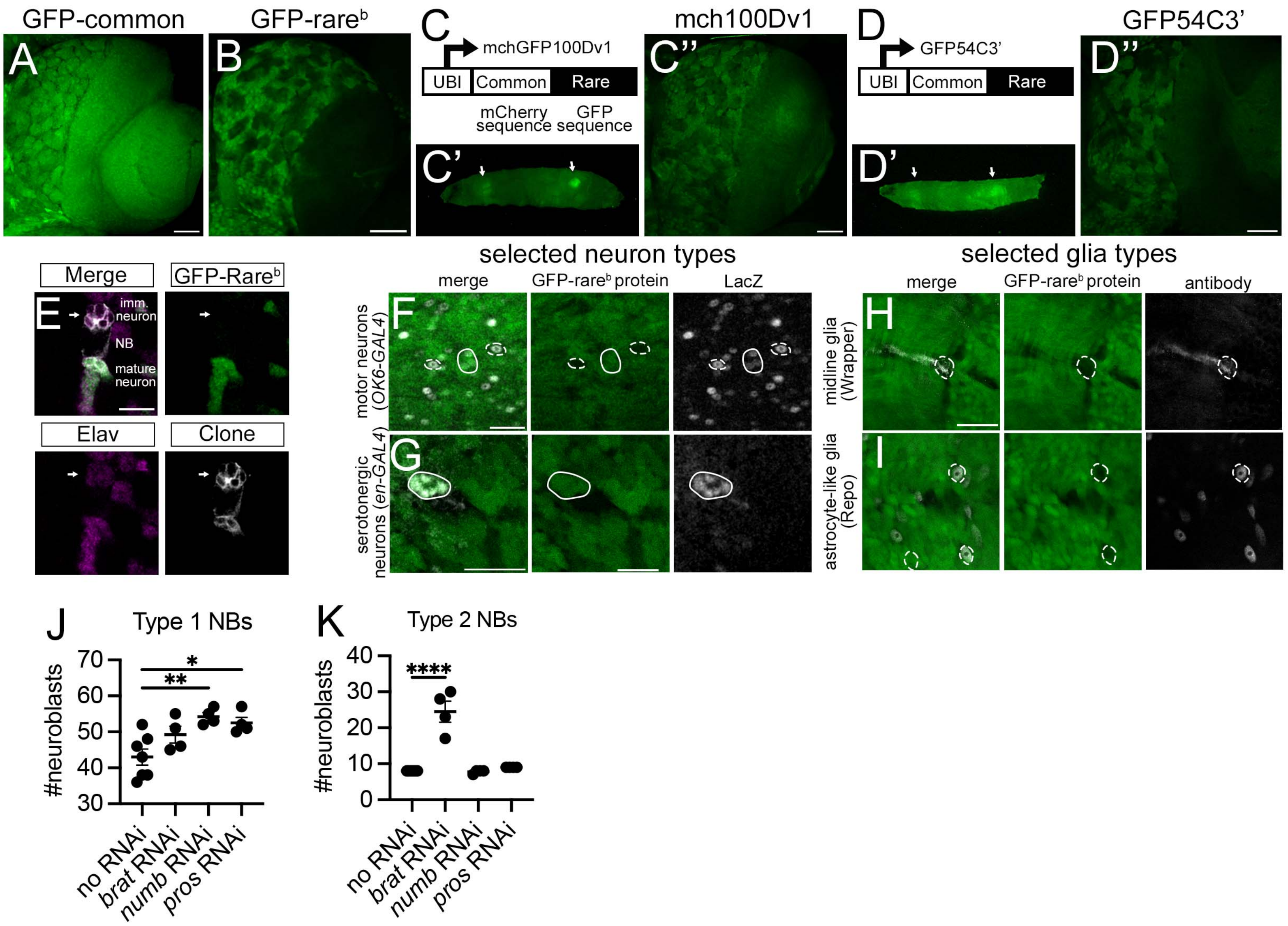

Figure S2: Orb2 whole-animal knockout and neuron specific knockdown result in similar decreases in GFP-rareb expression.

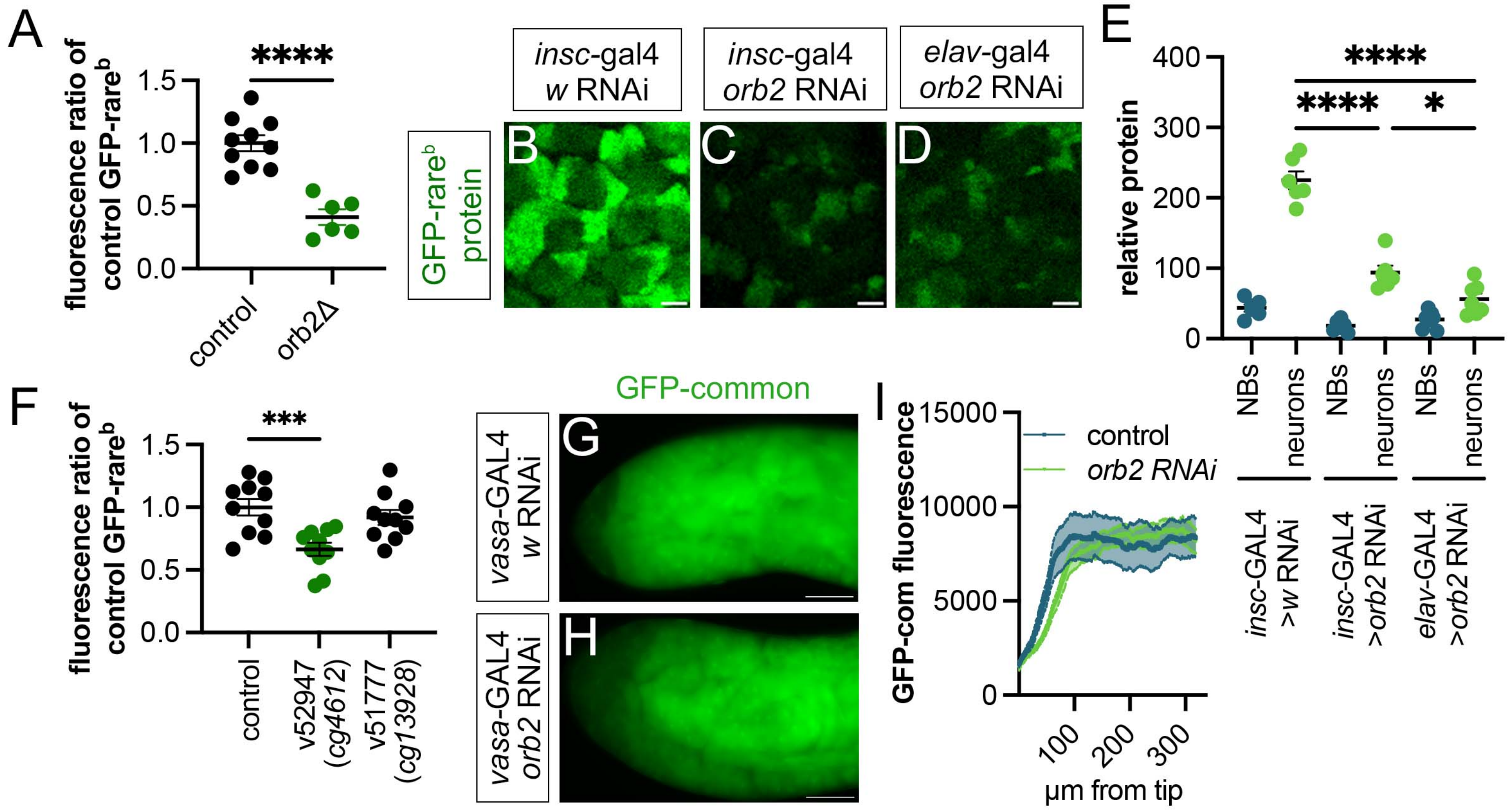

Figure S3: Both Orb2 binding sites and CAI are important to determine expression regulation in the Drosophila brain

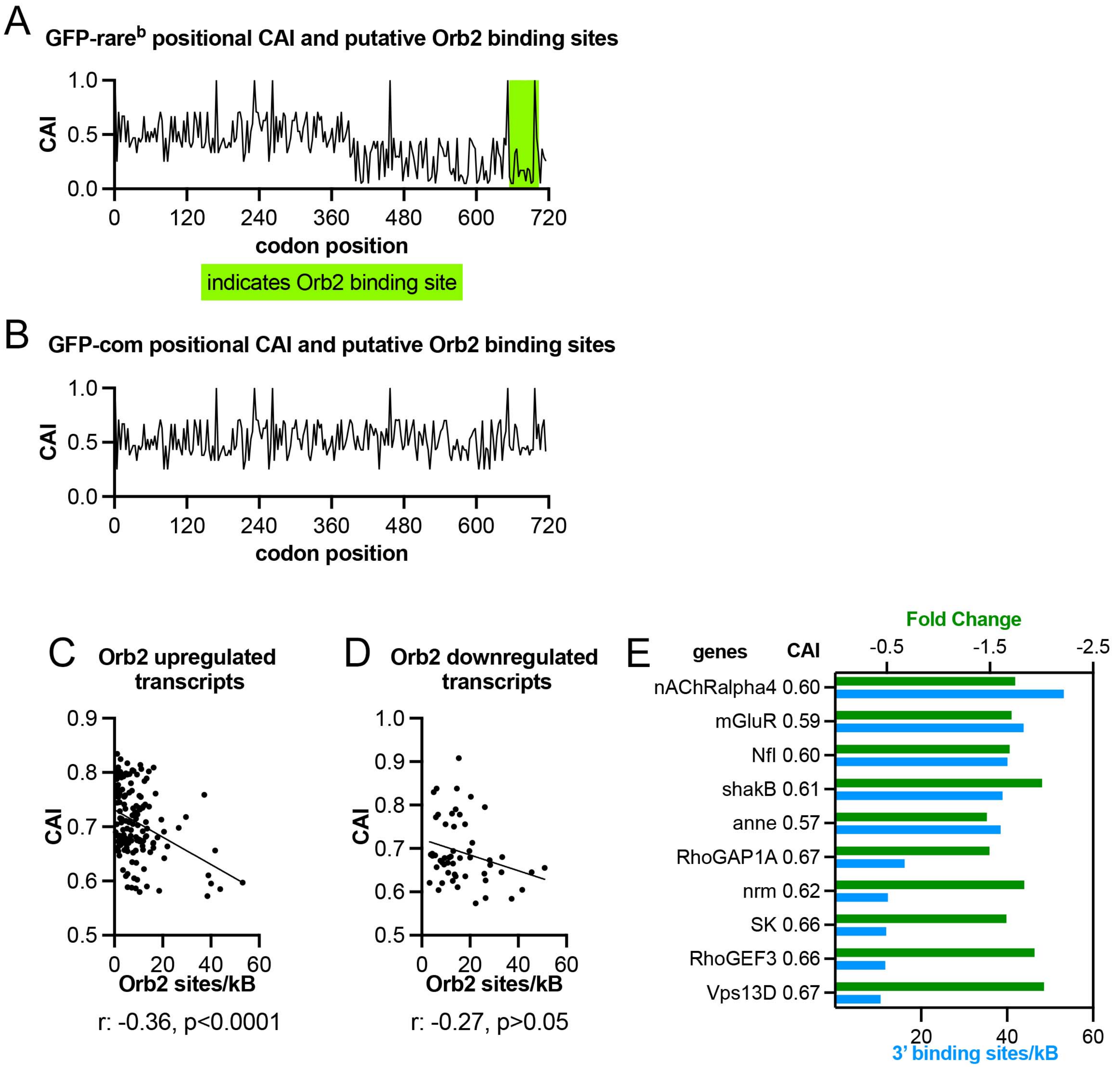
